# Genetic Fine-mapping with Dense Linkage Disequilibrium Blocks: genetics of nicotine dependence

**DOI:** 10.1101/2020.12.10.420216

**Authors:** Chen Mo, Zhenyao Ye, Kathryn Hatch, Yuan Zhang, Qiong Wu, Song Liu, Peter Kochunov, L. Elliot Hong, Tianzhou Ma, Shuo Chen

## Abstract

Fine-mapping is an analytical step to perform causal prioritization of the polymorphic variants on a trait-associated genomic region observed from genome-wide association studies (GWAS). The prioritization of causal variants can be challenging due to the linkage disequilibrium (LD) patterns among hundreds to thousands of polymorphisms associated with a trait. We propose a novel *ℓ*_0_ graph norm shrinkage algorithm to select causal variants from dense LD blocks consisting of highly correlated SNPs that may not be proximal or contiguous. We extract dense LD blocks and perform regression shrinkage to calculate a prioritization score to select a parsimonious set of causal variants. Our approach is computationally efficient and allows performing fine-mapping on thousands of polymorphisms. We demonstrate its application using a large UK Biobank (UKBB) sample related to nicotine addiction. Our results suggest that polymorphic variances in both neighboring and distant variants can be consolidated into dense blocks of highly correlated loci. Simulations were used to evaluate and compare the performance of our method and existing fine-mapping algorithms. The results demonstrated that our method outperformed comparable fine-mapping methods with increased sensitivity and reduced false-positive error rate regarding causal variant selection. The application of this method to smoking severity trait in UKBB sample replicated previously reported loci and suggested the causal prioritization of genetic effects on nicotine dependency.

**Author summary:** Disentangling the complex linkage disequilibrium (LD) pattern and selecting the underlying causal variants have been a long-term challenge for genetic fine-mapping. We find that the LD pattern within GWAS loci is intrinsically organized in delicate graph topological structures, which can be effectively learned by our novel *ℓ*_0_ graph norm shrinkage algorithm. The extracted LD graph structure is critical for causal variant selection. Moreover, our method is less constrained by the width of GWAS loci and thus can fine-map a massive number of correlated SNPs.

## 1 Introduction

Genome-wide association studies (GWAS) became the main tool to understand associations between genetic variants and complex biological traits such as nicotine addiction [1–4]. Many genetic loci have been discovered and replicated to be associated with nicotine and smoking traits [2,3,5,6]; however, even replicated loci may differ from the causal variants due to linkage disequilibrium (LD) among proximal and distant loci. Specifically, non-causal variants can be in association with a trait due to the high correlation with the causal variants due to LD [4,7,8]. Hence, the post-hoc examination of trait association results is required to select the likely causal variants from these that are associated with the trait due to LD effects. Fine–mapping algorithms can prioritize the likely causal variants by accounting for a complex LD structure and thus provide more insights into the underlying biological mechanism of the trait [4,8].

The variance of a polygenic trait can be associated with hundreds to thousands of SNPs, and a fine-mapping step aims to prioritize causal variants among all the genetic variants detected by GWAS [4]. From a statistical perspective, genetic fine-mapping is a variable selection procedure that identifies a parsimonious set of variants from a large number of correlated SNPs. Both regression shrinkage and Bayesian methods have been developed to implement genetic fine-mapping. Regression shrinkage methods such as least absolute shrinkage and selection operator (LASSO) and elastic net (ENET) have been applied; however, these methods suffer from a negative bias that leads to missed causal variants based on simulation studies [9,10]. Bayesian fine-mapping methods select causal variants by estimating the probability that an SNP is included as causal in the model according to the posterior inclusion probability (PIP) [11,12]. Yet, the computational cost of the Bayesian methods can grow exponentially with the length of the genomic region and, therefore, impractical for large sets (> 1000) [4,11,12].

An accurate fine-mapping model requires exact knowledge of the LD pattern among the SNPs within a GWAS locus [4,7,8,13,14]. Generally, the LD pattern refers to two aspects: i) the local aspect, pairwise LD scores between SNPs which are directly available, and ii) the global aspect, the (latent) network topological structure (e.g., the community structure) of the LD matrix. A large body of literature on multivariate statistics has shown that the accuracy of variable selection relies on the accurate knowledge of the network structure of the dependence between predictors [14–19]. However, the network structure of the LD pattern measured using the commonly used haplotype block methods cannot accurately reveal the latent network structure of the LD pattern. Specifically, the pairwise correlations among the SNPs within a haplotype block are not distinguishable from those outside blocks, which poorly characterizes the underlying LD network structure [20–22]. The obscure LD structure limits the accuracy of fine-mapping models [14]. To fill this gap, we propose a *ℓ*_0_ graph norm shrinkage approach to reorganize haplotype blocks into sets of dense LD blocks consisting of highly correlated neighboring SNPs. ‘Dense’ refers to the proportion of high LD scores (r^2^ > 0.8) between SNP pairs in the block is high (> 90%) [23]. Both the covariance matrix of SNPs and the network topology can be accurately estimated given the extracted dense LD block structure and then incorporated into the causal variant selection procedure. Here, we show that our approach improves the accuracy of fine-mapping versus approaches using haplotype block-based LD patterns (see details in subsection 2.1 and section 4).

We evaluated our fine-mapping approach with dense LD block using simulation studies and by replicating findings on the genetics of nicotine addiction using a large and inclusive UK biobank sample. The simulation results show that it can effectively improve the sensitivity and false-positive error rate in the causal variant selection by improving variable selection accuracy with dense LD structure [24–27]. We applied this approach in UKBB sample to focus on a gene cluster on chromosome 15 centered around IREB2, CHRNA3, CHRNA5, CHRNB4, HYKK, and PSMA4 genes that is associated with smoking severity measured as with a cigarettes per day – CPD trait [28–31]. Our fine-mapping analysis prioritized 93 variants with high statistical significance (*p* < 10^−50^), and most of them having large effect sizes (> 0.75). These causal variants reside across the genes in a highly correlated (average r^2^≈0.89) LD block. This leads to a systematic and multi-gene causal variant selection process, which may further our understanding of the underlying genetic mechanism for a complex trait.

## 2 Methods

### 2.1 Background and Motivation

#### 2.1.1 Background

We focus on a GWAS locus with a trait for a study cohort. Denote *X_n×p_* for the *p* SNPs in the GWAS locus for *n* participants, and *Y*_*n*×1_ for a scalar trait. We let ***R**_p×p_* represent the LD matrix, which can be obtained from the pre-calculated public depository or empirical estimation. The genetic fine-mapping aims to identify a small set of SNPs (e.g., *q < p* SNPs) associated with the trait while accounting for the ***R**_p×p_*. Since it is well-known that the variable selection procedure is influenced by the knowledge of the network structure of ***R**_p×p_*, we then demonstrate the rationale of choosing dense LD over traditional haplotype block.

#### 2.1.2 Motivation: dense LD block vs. haplotype block for causal variant selection

We start with demonstrating the input LD matrix ***R**_p×p_* in a GWAS locus from a data example in Fig 1(A) (see the details for data and materials in section 4). The LD matrix includes 1733 SNPs, which are correlated and exhibit an organized and latent network structure. Next, we show the revealed network structure by a haplotype block and dense LD block detection algorithms in Fig 1(B). Note, the detected LD network structure determines the order of SNPs in the dense LD structure. Clearly, the pairwise correlations between SNPs in a dense block are much higher than pairwise correlations between SNPs in a haplotype block. This is well-aligned with the fact that the LD score tends to increase between a pair of SNPs as their physical distance decreases, but the trend is neither linear nor monotone (see Fig 2). Therefore, it is sound to reorganize the order of SNPs based on their LD network structure regardless of their physical locations for the purpose of variable selection.

**Fig 1.**
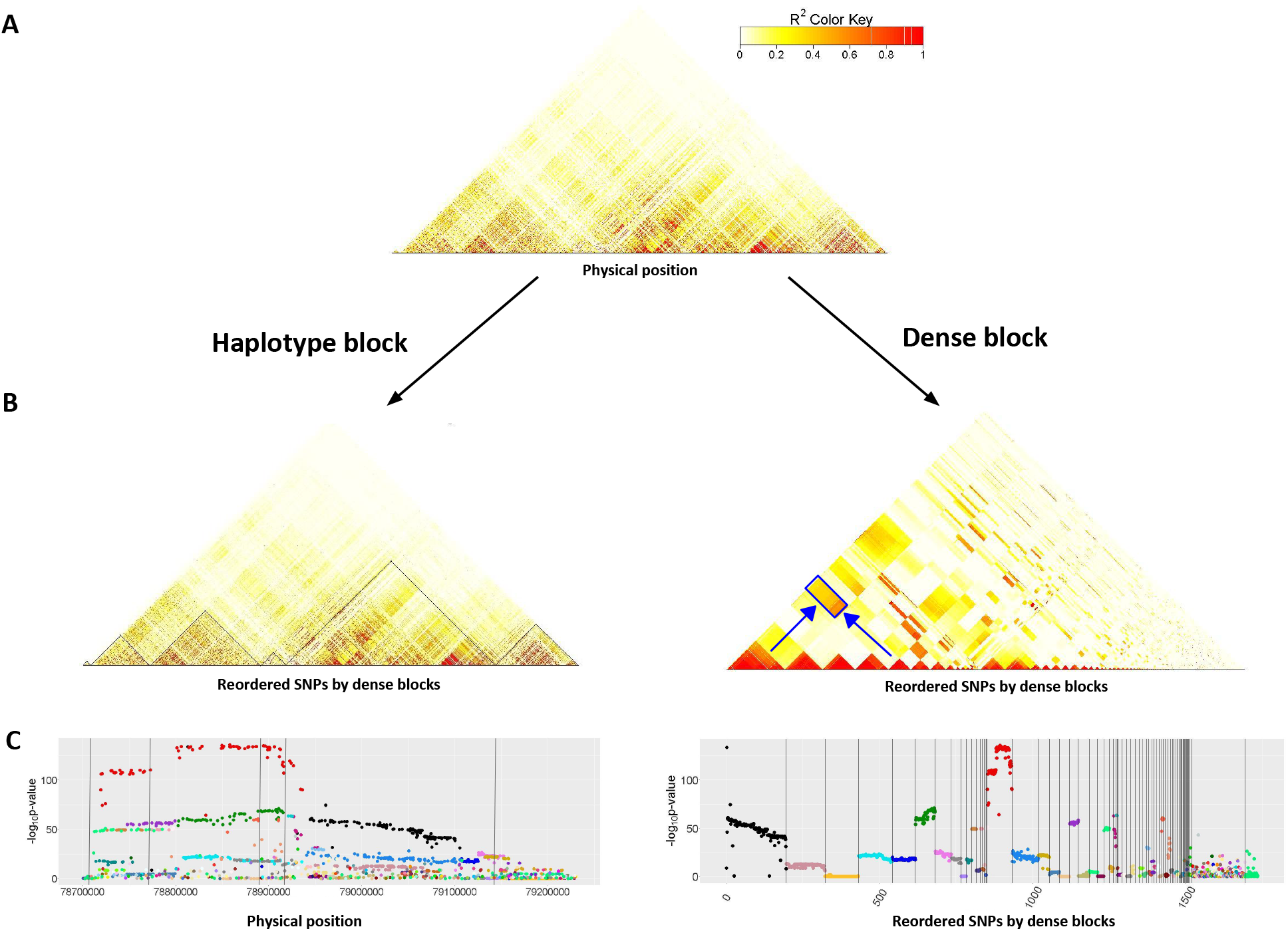
Manhattan plots and heatmaps before and after LD pattern detection via GSLD. (A) shows the raw heatmap of the genomic region. (B) shows the heatmaps of haplotype block (left) and dense block (right) learned from (A). (C) shows the Manhattan plots with SNPs ordered by haplotype block (left) and dense block (right). The vertical lines in the Manhattan plots indicate the block boundaries. The plots on the left list the SNPs in their natural physical position. On the contrary, the plots on the right reorder the SNPs based on rankings obtained from the LD pattern detection via *ℓ*_0_ graph norm shrinkage. SNPs are colored along with their block ID. The blue rectangle highlights an example of SNPs with distinct levels of trait associations, having moderate correlation (*r*^2^) ranging from 0.4 to 0.6, which are assigned to two different blocks.

**Fig 2.**
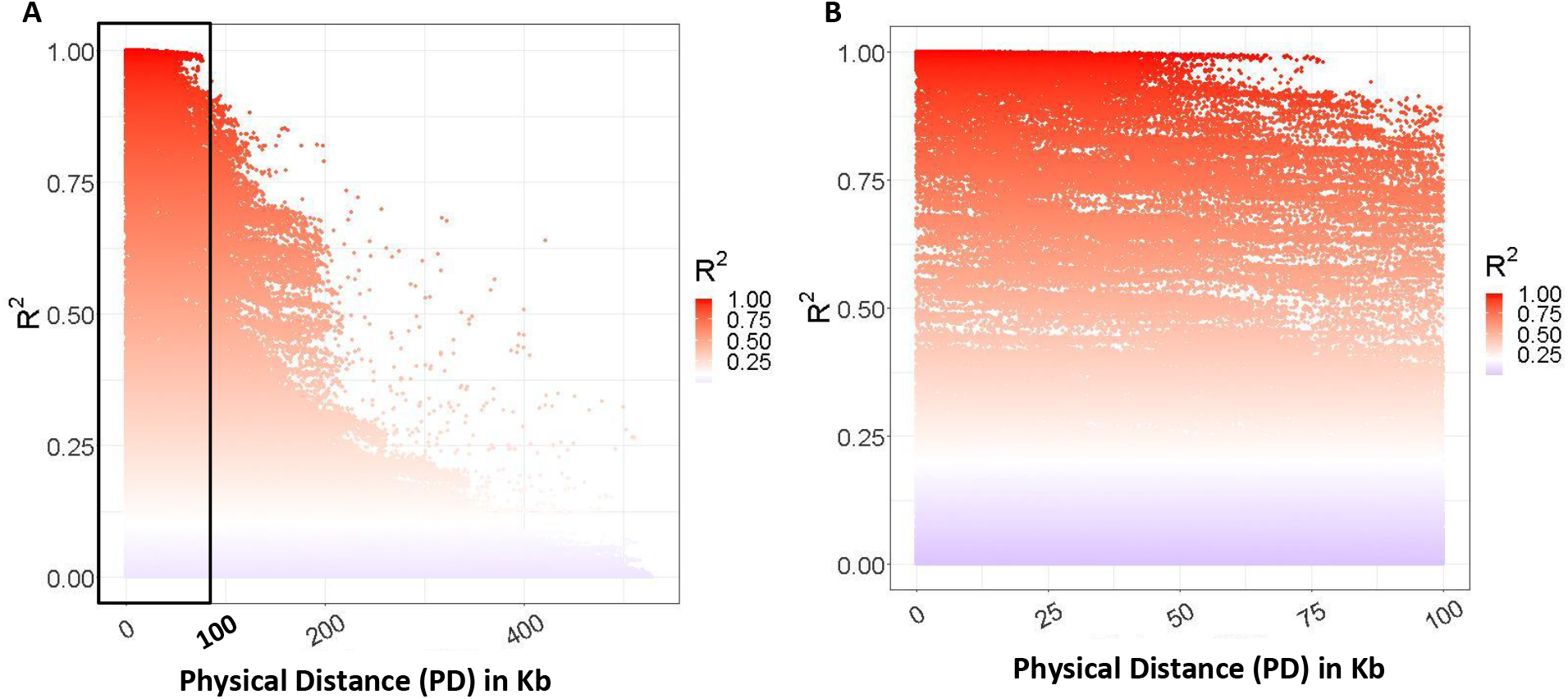
LD decay (in Kb) plot. A) displays the pairwise correlation (*r*^2^) of all SNPs pairs in the selected region from CPD data. The zoom plot in B) shows relationship of *r*^2^ and physical distance of pairs of SNPs within *PD* = 100 Kb and shows non-monotonic and nonlinear pattern.

**Fig 3.**
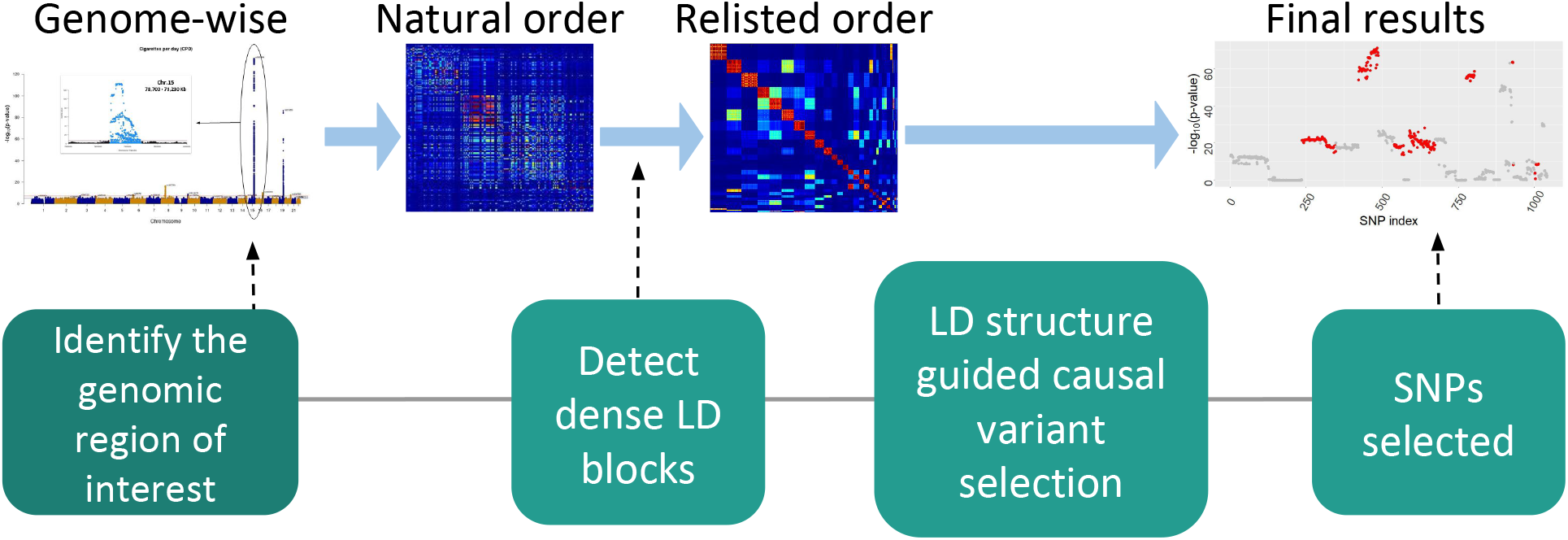
Overview of GSLD Procedures. A genomic region selected based on genome-wide analysis results is passed to GSLD fine-mapping following two steps: i) detect the LD structure of SNPs, and ii) select SNPs based on the SNPs data and the LD structure.

We further examine whether the LD pattern is related to the causal variant selection by jointly demonstrating the Manhattan plots and the LD pattern plots. In Fig 1(C), we note the statistical inference results of SNPs in a dense LD block are much more correlated than those in a haplotype block. For example, the red, blue, black colored SNPs are from three dense blocks. Although they are physically apart across multiple haplotype blocks, their statistical inference results are highly clustered. In other words, the variable selection can take into account the dense LD block structure to improve fine-mapping accuracy. We introduce our method to detect and exploit dense LD blocks for fine-mapping in the rest of this section and show the advantages of this strategy in sections 3 and 4. We also note that only the statistical inference results of SNPs in a dense LD block are coherent, while SNPs in a loosely correlated block show a large variety. Therefore, we are motivated to develop algorithms to detect dense LD blocks and the according fine-mapping procedure.

### 2.2 Detect dense LD blocks via *ℓ*_0_ graph norm shrinkage

Let **R** denote the LD score matrix for the wide genomic region, where each entry 0 ≤ *r_ij_* ≤ 1 represents the LD score between SNPs *i* and *j* (1 ≤ *i ≤ j ≤ n*). Let *G* = {*V, E*}be a graph notation for the LD pattern, where the vertice/node set *V* represents the *n* SNPs (|*V*| = *n*) in the genomic region of interest, and the edge set *E* indicates the levels of LD between the *n* alleles 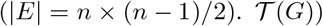 is the latent graph topology of the LD patterns which elucidates the assignment of nodes and edges into subgraphs {*G_c_*}(*G_c_* = {*V_c_, E + c*}) such that 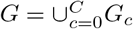. Specifically, we assume that *G* is composed by subgraphs {*G_c_*} of three categories: i) a set of dense LD blocks where all within block SNPs are highly correlated, ii) interactions between these blocks (i.e., edges with medium LD scores connecting dense blocks), and iii) the rest of the graph *G*_0_. We aim to extract the dense LD blocks which can further guide the causal variant selection.

Given the three categories of subgraphs in *G*, we let an entry *r_ij_* follow a marginal mixture distribution that 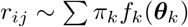, where *π_k_* is the proportion for a mixture component *k*(*k* = 1,⋯, *K*), and *f_k_*(***θ**_k_*) is corresponding distribution (e.g., beta distribution) with the parameters ***θ**_k_*. We assume that edges within dense blocks belong to a mixture component, interactions between blocks constitute *K* −2 mixture components, and edges from the rest of graph form a mixture component. We write the mixture distribution as:

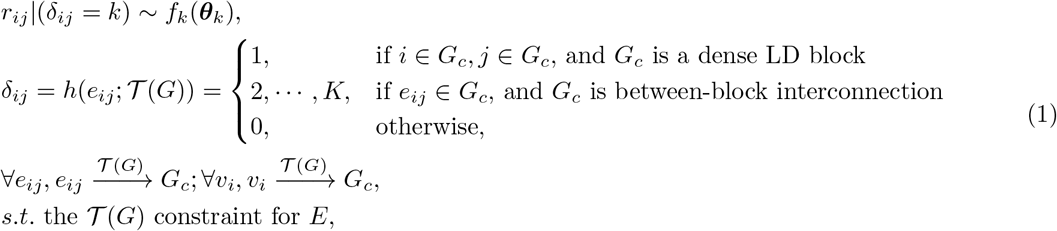

where *δ_ij_* is an indicator variable assigning an edge to a mixture component which is determined by 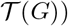) through a function *h*. We also have *E*(*r_ij_*|*δ_ij_* = 1) > *E*(*r_ij_|δ_ij_ = k, k* > 1) >*E*(*r_ij_*|*δ_ij_* = 0). The LD graph structure based mixture model is distinct from the conventional mixture model because we cannot freely assign an edge *r_ij_* to any mixture component. For example, given a dense LD block *G_c_*, we have {*e_ij_ ∈ G_c_* and *e_ik_ ∈ G_c_*} ⇒ *e_jk_ ∈ G_c_*. Therefore, the estimating methods including EM and nonparametric Bayes models may not be directly applicable.

We consider the latent graph topological structure 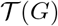 as the key parameter of the LD graph structure based mixture model because estimating ***θ**_k_*(*k* > 2) is straightforward given 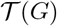. Although *K* is unknown in our infinite mixture model, it can also be estimated by the likelihood principle with a given 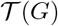. For the purpose of fine mapping, our primary goal is to detect dense LD blocks while prohibiting false-positive edges (with medium LD scores) being included in the blocks.

To link the underlying dense LD blocks with the input data **R**_*n×n*_, we introduce a matrix **U** = {*u_ij_*}_1≤*i,j≤n*_ obtained by thresholding **R**_*n×n*_:

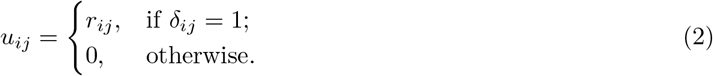

Then, our new objective function is:

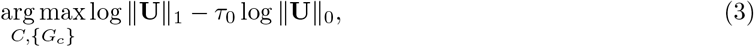

where 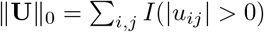 is the element-wise *ℓ*_0_ matrix norm (i.e., graph (size) norm), 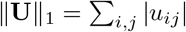 is the element-wise *ℓ*_1_ matrix norm, and 0 < *τ*_0_ < 1 is a tuning parameter. We maximize ||**U**||_1_ to assign a maximal number of high LD score edges into dense blocks with high sensitivity. In the meanwhile, we penalize the *ℓ*_0_graph norm ||**U**||_0_ to prohibit including false-positive and medium LD score edges into blocks. For example, assigning one false-positive SNP into a dense block comes with a high of greatly increasing the ||**U**||_0_ term, which is against our objective function. Note that *δ_ij_* = 1, only if *e_ij_ ∈ G_c_* and *G_c_* is a dense LD block. Therefore, we regulate the size of each dense block to ensure the low false-positive rate. The tuning parameter *τ*_0_ controls the level of parsimony. In general, larger *τ*_0_ leads to more dense yet smaller sized LD blocks. In practice, we can optimize *τ*_0_ based on the likelihood function of the mixture model as follows:

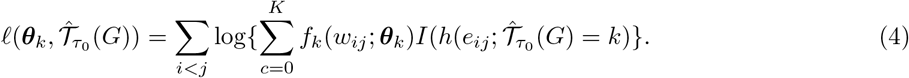

By implementing the *ℓ*_0_ graph norm shrinkage, we can accurately recognize the underlying dense LD blocks which can guide our causal variant selection. In practice, the direct optimization of Eq 3 is NP (nondeterministic polynomial time) complex and resort to the computationally affordable algorithm (see the appendix for details).

### 2.3 Causal variant selection based on dense LD blocks

Penalized regression is one of the popular approaches for fine-mapping, which jointly analyze multiple variants in a region. However, popular variable selection models like LASSO and ENET tend to randomly select single variants when those variants are highly correlated with adjacent SNPs [24,25,27,32]. Recently, fused-LASSO, a structured regularization method, has been developed to encourage similarity within a group of variables while retaining the overall sparsity [24,25,27,32]. This method naturally suitable for fine-mapping for the purpose of utilizing dense LD blocks in casual variant selection. Our newly developed method, GSLD, is built on top of the advances in penalized regression shrinkage, accounting for the structural patterns found in LD matrix. Based on the estimated dense LD block obtained according to Eq 3 and Eq 4, we then use the graph guided fused-LASSO, integrating the objective function as:

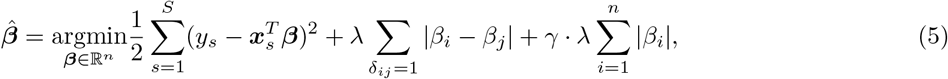

where *Y* ={*y*_1_, *y*_2_,…, *y_S_*}, *s* = 1, 2,…, *S* are the phenotype for *S* subjects in our data, ***x**_s_* = {*x*_*s*,1_, *x*_*s*,2_,…, *x_s,n_*}^*T*^(0, 1, 2), *i < j* = 1, 2,…, *n* are the *n* genotypes of the *s^th^* subject, and ***β*** = {*β*_1_,⋯, *β_n_*} represents the phenotype-genotype associations. The graph object converted from adjacency matrix was passed as term |*β_i_ − β_j_*| for regulation. *λ* and *γ* are the regulation parameters that control the weight of penalty terms in Eq 5. The regulation becomes a pure fusion of the coefficient vector beta when *γ* = 0; otherwise, a non-zero *γ* introduces a ratio of sparsity to terms |*β_i_*| and |*β_i_−β_j_*|, corresponding to the sparsity of coefficients and the sparsity of their differences, respectively. Given the difference between coefficients, fused-LASSO does not only provide shrinkage on single coefficients but also fuses some consecutive variables to have equal coefficients with each other; therefore, fused-LASSO imposes a similarity between correlated features and achieves a group-wise selection of variables. Further details for implementing the objective function can be found elsewhere [24,25,27].

In summary, GSLD can select causal variants based on the detected LD patterns. Since the *ℓ*_0_ graph norm shrinkage is applicable for a broad genomic region, GSLD can evaluate thousands of SNPs by treating them as a single fine-mapping region. This advantage provides universal consideration for all SNPs in the entire genomic region, accounting for their correlations. Accordingly, this approach can improve the causal variant selection in many applications. We demonstrate the performance of our method using a data example and extensive simulations.

## 3 Results

We applied our method to UK Biobank (UKBB) data to investigate the genetic factors for nicotine addiction [33].

### 3.1 UKBB data

UKBB database is an international resource that provides abundant health information, including genetic data of 500,000 volunteer participants [33]. We focus on investigating the association between genetic factors and nicotine addiction by GWAS, using cigarettes per day (CPD). CPD is one of the traits that have been used to evaluate the phenotype-genotype associations for nicotine addiction and smoking behavior [18,33], and it was defined using three data fields from UKBB (see S1 Appendix for the detailed definition). We restricted our study cohort to Caucasian current and previous smokers that consists of 142,752 participants. The age of the participants ranged from 40 to 72, involving 74,061 males and 68,691 females. GWAS was first carried out to identify a genomic region of interest, which contains SNPs having strong significant phenotype-genotype associations and requires fine-mapping for extensive causal variant selection. The results showed that a genomic region on chromosome 15 (15:78,700,000 to 79,230,000) had the strongest signal for CPD (see S1 Appendix for the genome-wide Manhattan plot). These loci coincided with the well-known cluster of genes IREB2, CHRNA3, CHRNA5, CHRNB4, HYKK, and PSMA4 that have been discussed in several GWASs [28–31]. To further identify causal variants, we applied our fine-mapping approach for an in-depth evaluation of this genomic region, covering 5.3 Mb and 1733 SNPs.

### 3.2 Results by Dense LD Block

We implemented the GSLD to assess the effect of 1733 SNPs on CPD by following the two-step procedure. As shown in Fig 4, GSLD detected 68 dense blocks based on the LD matrix obtained via PLINK of the selected genomic region [34]. The block detection performance of GSLD was stable since the block structure is almost invariant to the tuning parameter. While integrating the LD structure characterizing by dense LD blocks, GSLD selected 93 SNPs for CPD through analyzing the joint effect of multiple SNPs using graph-guided regression shrinkage. The results are visually presented in Fig 4. Owing to the cluster-wise penalty term (Eq 5), we identified 3 clusters (i.e., the 18^*th*^, 37^*th*^, and 41^*st*^ blocks) of causal variants instead of individual SNPs. Among the 93 chosen SNPs, 81 SNPs are from the 18^*th*^ block. The 37^*th*^ and 41^*st*^ blocks have 10 and 2 SNPs, respectively. Although the 12 SNPs are partitioned into different blocks, some of them are from genes IREB2 and MORF4L1, which are functionally related to smoking-associated diseases and behaviors [29,35]. Additionally, 78 of the 93 SNPs have published gene information [36,37]. Many of them are located in the well-known gene cluster related to smoking-associated diseases and behaviors [30]. These SNPs are shown in different colors in Fig 4. Among the 15 SNPs without published gene information, 3 SNPs (1 in the 18^*th*^ block and 2 in the 37^*th*^ block) have been discovered as associated SNPs for smoking cessation and chronic obstructive pulmonary disease (COPD) [38–40]. These findings are encouraging as similar results of these genes have been replicated across studies, including some investigations based on UKBB data [7,28–31,41,42]. Further investigations are needed to discover the potential hidden functional mechanisms for the associations between smoking and the SNPs without published knowledge.

**Fig 4.**
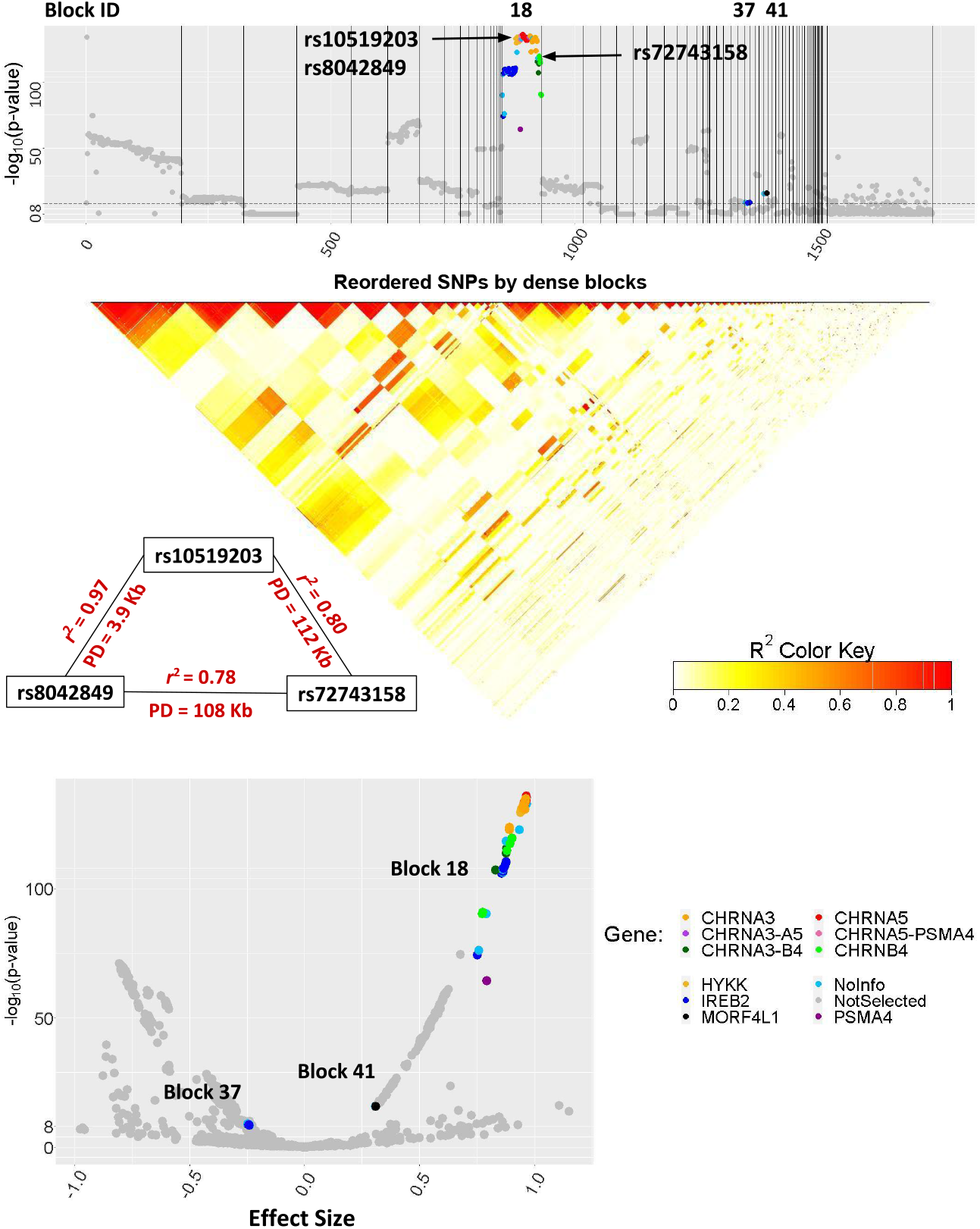
Results of GSLD for CPD. The Manhattan plot shows the SNPs excluded in grey and highlights the selected SNPs in other colors. The SNPs are colored according to the genes they are located in. The horizontal dashed line (at −log_10_(p-value) = 8) corresponds to the commonly used genome-wide significance level (p-value = 5 × 10^−8^). The block IDs of selected SNPs are shown on top of the plot. The volcano plot shows that SNPs in the selected blocks, 18^*th*^, 37^*th*^, and 41^*st*^, are gathered relatively closed, indicating similar trait associations within a block regarding p-values and effect sizes. The arrows point out three SNPs (rs10519203 (in HYKK), rs8042849 (in HYKK), and rs72743158 (in CHRNB4)) with strong correlations. One of them has large physical distances with the others whilst their correlations are strong (rs72743158 and rs10519203: *r*^2^ = 0.80, *PD* = 112*Kb*; rs72743158 and rs8042849: *r*^2^ = 0.78, *PD* = 108*Kb*; rs10519203 and rs8042849: *r*^2^ = 0.97, *PD* = 3.9*Kb*), indicating that high LD can occur in both neighboring and distant loci.

Moreover, we found that the SNPs in the same block are highly correlated with each other, although they come from different genes. In some cases, high LD could be observed between SNPs though they are physically distant. An example that well demonstrates the high LD between physically distant SNPs were identified in the selected 18th block (see Fig 4). Three SNPs on chromosome 15 (rs10519203 (in HYKK), rs8042849 (in HYKK), and rs72743158 (in CHRNB4)) have high LD scores (*r*^2^ ⩾ 0.78) with each other, yet one of them has a physical distance (PD) greater than 100 Kb with the others. Previous work suggested that these SNPs might harbor shared causal variants that explain the local GWAS and expression quantitative trait loci (eQTL) signals associated with COPD [43]. They may jointly affect the expression of candidate COPD genes; nonetheless, given their physical positions, they could be placed into different haplotype blocks using existing methods. On the contrary, our *ℓ*_0_ graph norm shrinkage algorithm captured their pairwise correlations and assigned them to the same dense LD block despite the large distance.

Interestingly, aside from the strong pairwise correlation, SNPs within a dense LD block have similar effect sizes and statistical significance (see the volcano plot in Fig 4), which can be explained both mathematically and biologically. Given the dense blocks, these variants potentially would inherit together due to evolutionary forces, such as natural selection, recombination, and genetic drift. And they are likely related to each other concerning biological functions. Additionally, we applied other fine-mapping methods, including LASSO and ENET, for CPD [9,10]. Bayesian models are not used because this genomic region is overwhelmingly vast and requires extensive computational capability. The fine-mapping results of LASSO and ENET are summarized in Fig 5. LASSO and ENET only identified 14 causal variants since they did not utilize LD patterns in the variable selection algorithms, yielding lower accuracy, especially sensitivity. The causal variants were sparsely distributed in the studied genomic region.

**Fig 5.**
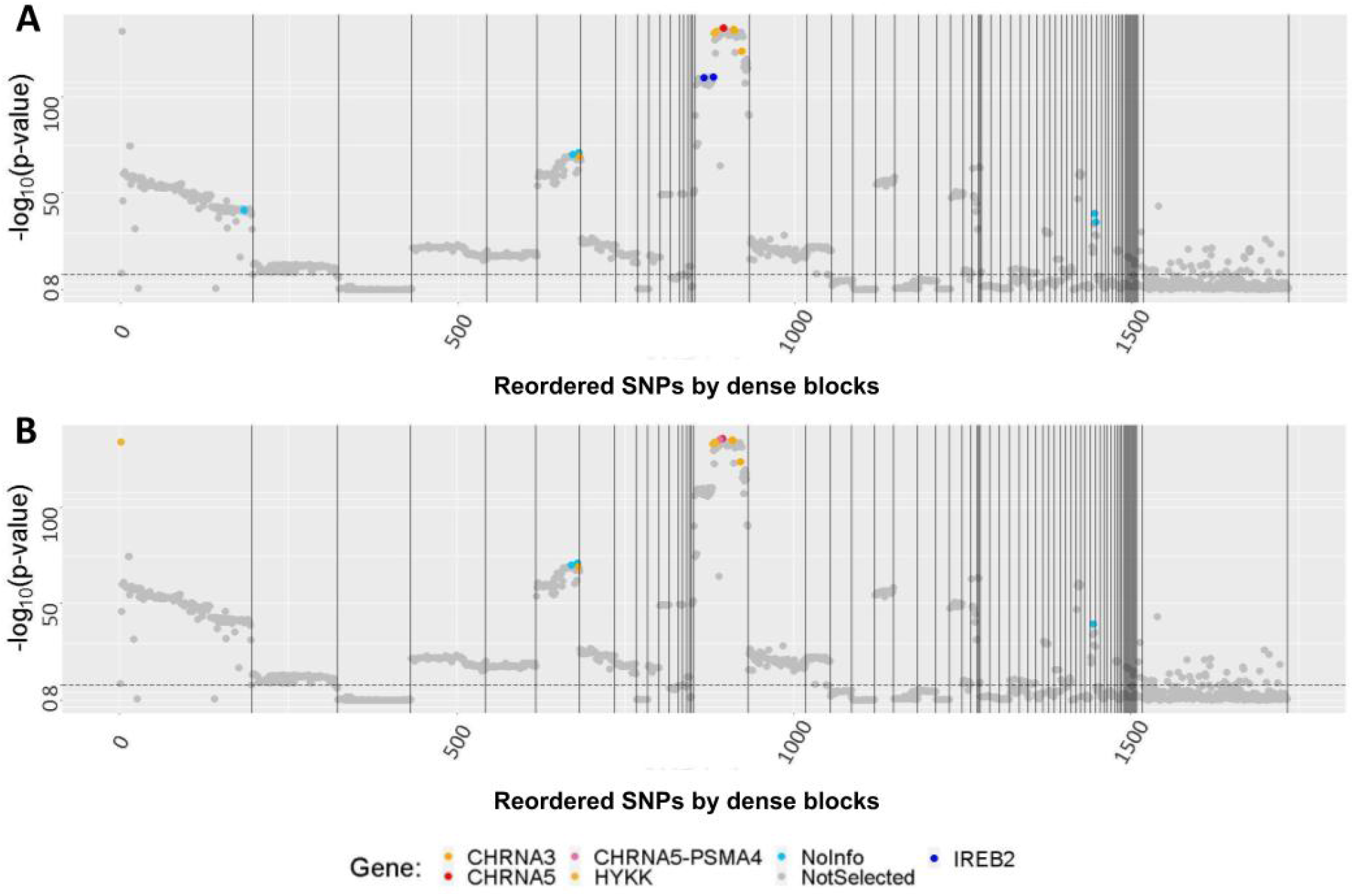
Selected SNPs for CPD via LASSO and ENET. The Manhattan plots show the SNPs excluded in grey and highlight the selected SNPs in other colors. A and B reveal the results of LASSO and ENET, respectively. SNPs are aligned in the relisted order from GSLD. The horizontal dashed line (at −log_10_ (p-value) = 8) corresponds to the genome-wide significance level (p-value = 5 ×10^−8^).

## 4 Simulations

### 4.1 Synthetic Data

We implemented simulation studies to evaluate the performance of GSLD. We first generated genotype data using the SNP simulation tool HAPGEN2, mimicking the LD pattern as the HapMap3 and 1000 Genomes Project [44]. The LD matrix was then acquired using PLINK based on the simulated genotypes [45]. Specifically, we focused on a genomic region including 1000 SNPs on chromosome 15, with the physical position starting from 78,700,000 to 79,230,000 bp. For a subject s, we denote the simulated vector of genotype data as ***x**_s_*, where *s*= 1,⋯, *S* and *S* = 500. Next, we simulated phenotype data based on the genotype data and causal variant related parameters based on a regression model:

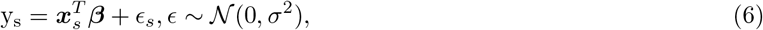

where *y_s_* is phenotype, ***β***_*n*×1_ is a vector of parameters reflecting the genotype-phenotype association, and *ϵ_s_* is the residual following a normal distribution with a zero mean and variance *σ*^2^. We consider ***β***_*n*×1_ as our primary parameters of interest since our goal is to accurately recover the true phenotype-genotype association 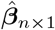 based on *Y* and ***x***. In light of the patterns of genotype-phenotype associations from empirical GWASs, we set the causal variants based on the dense LD blocks. Let the parameter for SNP *i β_i_* = *b|i* ∈ *G*_1_ or *G*_2_ where *G*_1_ and *G*_2_ are two dense LD blocks and *b* is the effect size, and *β_i_* = 0 otherwise. We set three settings of the effect size *b* = 0.6, 0.8, 1 and repeated the above procedure 100 times for each setting.

In the simulation, the tuning parameters were selected based on sensitivity and specificity across 100 repeated analyses. For comparison, we also applied the commonly used fine-mapping models based on penalized regression shrinkage methods (LASSO and ENET) and Bayesian methods (PAINTOR and JAM) [46–49]. We exploited the maximum computational capacity (i.e., using the maximum length of the genomic region and number of causal variant candidates) to select causal variants via Bayesian methods. Note that the width of the genomic region and the number of causal variants may be constrained in many Bayesian fine-mapping models. The performance of GSLD and the comparable methods was evaluated at the end of the simulation.

#### 4.1.1 Benchmark and evaluation criteria

We evaluated the performance of GSLD and comparable methods based on the accuracy of causal variant selection, and the results are summarized in Table 1. Specifically, the sensitivity and specificity are calculated as

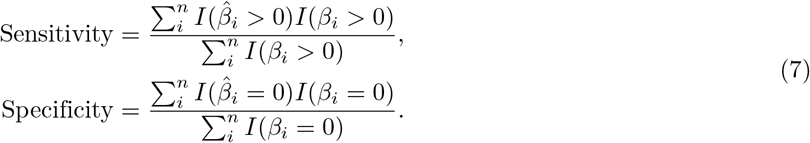

**Table 1.** Summary of performance for various methods in the simulation.

| Beta |  | GSLD | LASSO | ENET | PAINTOR | JAM* |
| --- | --- | --- | --- | --- | --- | --- |
| **0.6** | *Sensitivity* | 1.0000(0.0000) | 0.7607(0.0058) | 0.8128(0.0055) | 0.1012(0.0011) | 0.5332(0.0051) |
|  | *Specificity* | 0.9563(0.0009) | 0.9167(0.0029) | 0.9152(0.0022) | 0.9151(0.0002) | 0.8175(0.0048) |
| **0.8** | *Sensitivity* | 1.0000(0.0000) | 0.6826(0.0065) | 0.7340(0.0053) | 0.0978(0.0002) | 0.4380(0.0052) |
|  | *Specificity* | 0.9561(0.0003) | 0.8693(0.0040) | 0.8576(0.0029) | 0.9155(0.0002) | 0.8498(0.0032) |
| **1** | *Sensitivity* | 1.0000(0.0000) | 0.7991(0.0051) | 0.8388(0.0047) | 0.0983(0.0004) | 0.4203(0.0065) |
|  | *Specificity* | 0.9485(0.0003) | 0.8891(0.0028) | 0.8689(0.0023) | 0.9158(0.0002) | 0.8592(0.0035) |

The table reports mean (standard error) of 100 simulation.
JAM\*: 243 SNPs are used to calculate the sensitivity and specificity calculation here due to SNP size limitation.

### 4.2 Simulation analysis results

Motivated by our data example and many GWASs, we consider all SNPs in a dense LD block as causal variants, rather than selecting a few causal variants and treating others as correlated SNPs. Under this setting, our method outperforms the comparable methods regarding sensitivity and specificity. The sensitivity of GSLD is higher than competing methods because it is built on the accurately learned LD graph topological patterns and selects causal variants systematically. The sensitivity of Bayesian methods are lower due to the constraint for the number of causal variants. When the genomic region is wide, regression shrinkage models seem more suited.

## 5 Discussion

We have developed a novel fine-mapping method to identify causal variants while incorporating the LD structure based on dense LD blocks for a genomic region. This method innovatively characterizes the complex LD matrix by detecting blocks with condensed correlations to assist causal variant selection. When high LD occurs between distant SNPs, our method becomes exceptionally beneficial as it can fine-map the whole region without separating it into subregions. Overall, when assessing up to thousands of SNPs, our method offers comprehensive genomic information in mapping causal variants while maintaining efficient computation.

Our fine-mapping approach is the first to explore the latent structure of the input LD matrix fully. In order to obtain the hidden information, we carry out detection based on the *ℓ*_0_ graph norm shrinkage to discover dense LD blocks. This algorithm is fundamentally different from traditional clustering and the community detection method used in haplotype block detection. The difference is due to the LD detection in GSLD that only assigns highly correlated SNPs into the same block without following their physical order. In contrast, existing methods often partition SNPs into blocks following the physical order yet cannot guarantee the denseness of their matrices. Our results demonstrate the importance of detecting groups of SNPs having strong correlations, showing its critical role in the fine-mapping analysis. In other words, detecting the dense blocks is analogous to discover a set of SNPs having similar effect sizes and statistical significance. Based on the dense LD blocks, GSLD tends to select a set of causal variants block-wise as it aggregates SNPs with similar trait associations (e.g., chosen SNPs for CPD are mainly from the 18^*th*^ block) but may not be physically closed. Hence, these SNPs could come from different genes. As mentioned above, GSLD can hardly discriminate a few causal variants from others in the same block, treating them as non-causal variants. In this situation, the classification between causal and non-causal variants can be further made with additional information, such as utilizing the knowledge from eQTL. Prior biological knowledge of their functional annotation (e.g., ENCODE) and downstream regulatory mechanism (e.g., eQTL) can be incorporated to prioritize causal variants and distinguish them from non-causal variants. Alternatively, the Bayesian fine-mapping approaches can be applied to each dense block for causal variant selection based on the posterior inclusion criteria.

By contrast with GSLD, traditional penalized regression shrinkage methods (e.g., LASSO and ENET) tend to select a few SNPs. It is unsurprising since they naturally shrink most coefficients among highly correlated variables, leaving a few SNPs with strong marginal trait associations. Both the application and simulation results exhibit the outstanding performance of GSLD in fine-mapping of a genomic region covering thousands of SNPs, comparing to LASSO and ENET. Moreover, due to the exhausting computational challenge, we could implement fine-mapping in a limited number of SNPs and consequently selected fewer SNPs via Bayesian. Although SNPs can be partitioned into different groups based on their LD patterns and reduce the dimension of data and computation time, the fine-mapping would only be implemented localized within a subregion. Accordingly, the model loses the comprehensive information across genes of the whole mapping region, ignoring the strong correlations among distant SNPs. Nevertheless, Bayesian fine-mapping would be efficient when the studied region is narrower. Therefore, when one needs to fine-map the entire genomic region, it is difficult to successfully select causal variants via traditional regression shrinkage and Bayesian fine-mapping methods.

To the best of our knowledge, our paper is the first to demonstrate the intrinsic LD patterns in dense blocks, and more importantly, that SNPs in a dense block can come from different genes yet be uniformly associated with the phenotype. In our approach, we want to emphasize that the SNPs assigned to the same block all have strong correlations; though, they are not necessarily physically contiguous (see plots of dense block in Fig 1). In future research, we plan to investigate biological knowledge about the functional links across genes potentially related to human traits.

In conclusion, we provide a new fine-mapping toolkit for a genomic region with a wide range. Based on the proposed objective function (Eq 3), our *ℓ*_0_ graph norm shrinkage method can effectively capture the latent structure and characterize the LD matrix by dense LD blocks for a genomic region. The LD structure coincides with the pattern of associations between SNPs and a phenotype; therefore, the dense structure can guide causal variant selection. Our simulation studies have demonstrated improved performance of GSLD regarding the accuracy of causal variant selection. The UKBB nicotine addiction study further shows that our method can extract causal variants in dense blocks with large effect sizes and small p-values. We also notice that the computational cost of GSLD is modest, and it can be applied to a variety of phenotypes (e.g., complex diseases) simultaneously. Therefore, the recent advances in graph-guided statistical models can beneficially assist in improving fine-mapping in genetic research.

## Supporting Materials

## S1 Appendix. Information of cigarettes per day (CPD) from UK Biobank

CPD was defined as the average number of cigarettes smoked per day by participants who were either current or past smokers and combined the data from UKBB fields 2887 (number of cigarettes previously smoked daily), 3456 (number of cigarettes currently smoked daily), and 6183 (number of cigarettes previously smoked daily (current cigar/pipe smokers) [33]. We recoded values smaller than 1 to 0 and constrained extreme values larger than 60 to 60; otherwise, we kept the original values as reported in UKBB. The final analysis included 142,752 participants aged from 40 to 72, involving 74,061 males and 68,691 females who had genetic data available in UKBB.

**S1 Fig.**
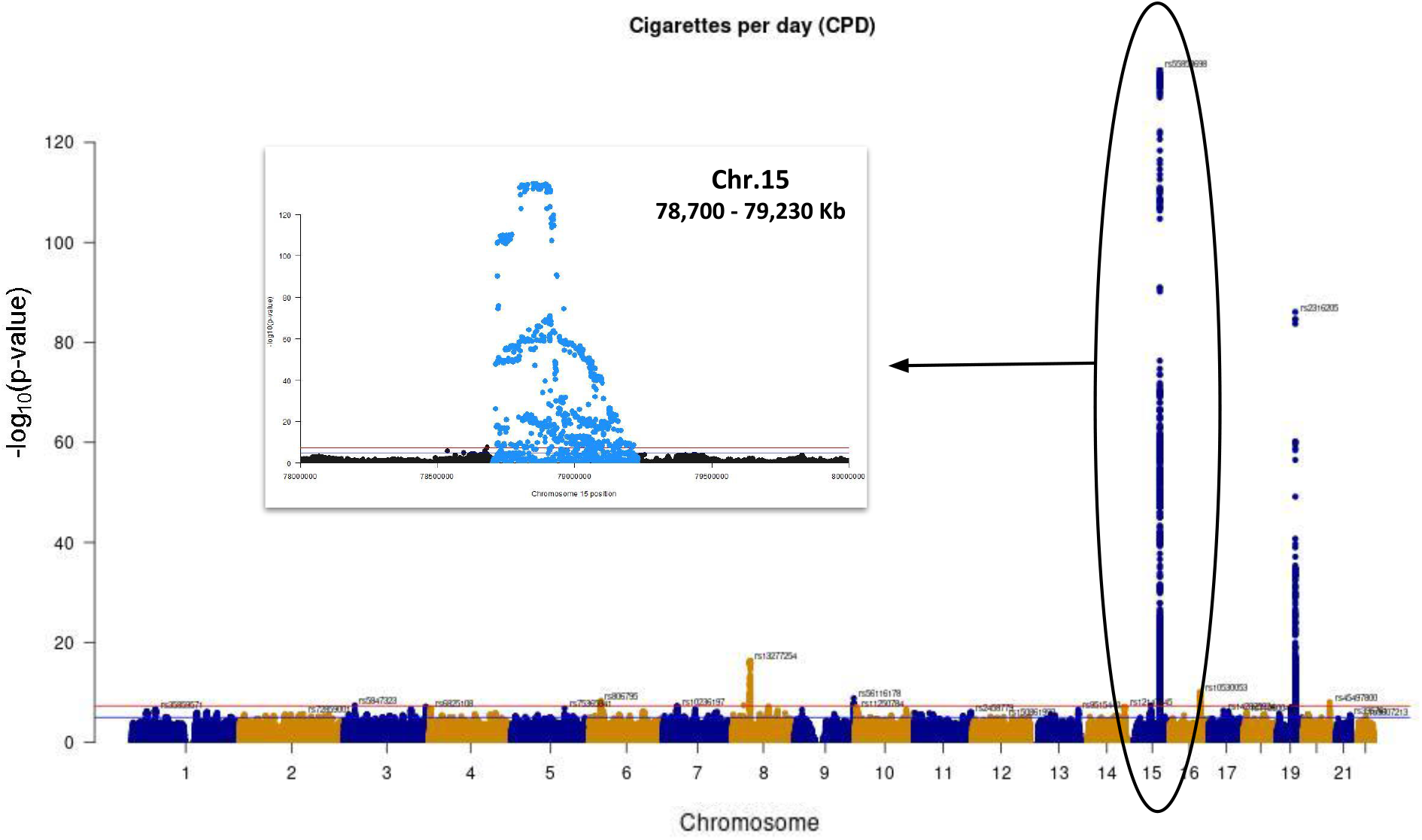
Genome-wide Manhattan plot and the genomic region of interest for cigarettes per day (CPD) Genome-wide analysis was performed to evaluate the associations between SNPs and CPD. The inclusive criteria, involving minor allele frequency (MAF) > 0.01, Hardy-Weinberg equilibrium (HWE) > 0.001, missingness per marker (GENO) > 0.05, and missingness per individual (MIND) > 0.02, were used for filtration in GWAS. We identified a region with extraordinarily strong association in chromosome 15 (circled in the Manhattan plot). In the zoom plot of chromosome 15, we see that the SNPs colored in light blue have extremely high −log_10_(p-value) between 78,700,000 and 79,230,000 bp. Therefore, we further explored the potential causal variants of the 1733 SNPs within this region and CPD via GSLD and compared the selection performance with other fine-mapping methods.

## S2 Appendix. Optimization of the objective function

The optimization of the objective function (1) is implemented by exhaustive search for *C* and estimating 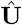 at each *C*. With a given *C*,

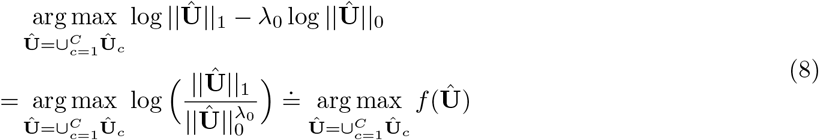

By default λ_0_ = 0.5 reflects balanced covering quality and quantity of true positive edges, and the objective function 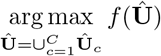 then becomes the well-known problem of *k* dense subgraph discovery, where *f*(·) is the density function. The problem has been solved in polynomial time by Goldberg’s min-cut algorithm [50] and a greedy algorithm with 1/2 approximation by [51]. In addition, the default topological community structure can be considered as quasi-cliques and the problem can be solved by additive approximation algorithms and local-search heuristics [52]. Alternatively, with the mild spatially-invariant assumptions that 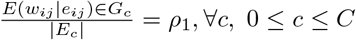, and 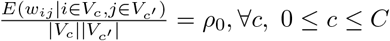 the primary objective function is equivalent to

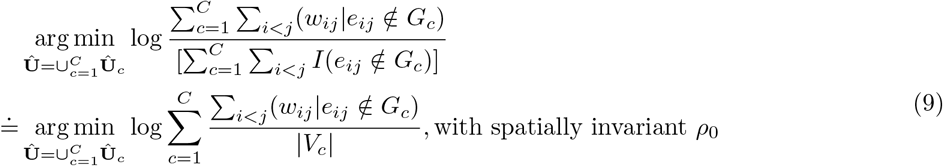

Although the objective function (9) is not convex, the issue of local optima in the discrete optimization can be solved by restarting the algorithm several times with different initialization and/or through orthonormal transforms [53, 54]. The proposed algorithm may better extract multiple weighted dense subgraphs (with an unknown number and unknown sizes of dense subgraphs) than the existing algorithms of dense subgraph discovery [34]. We then choose the optimal *C*\* by grid searching that maximizes the following criteria:

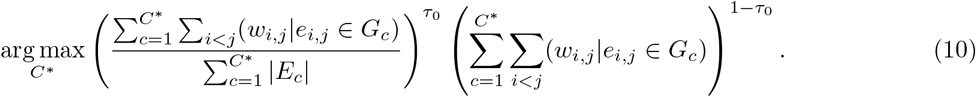

The criteria (10) can be directly derived from our primary objective function that 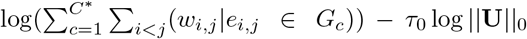. The first term in (10) indicates the ‘quality’ (the area density) of the extracted subgraphs, while the second term represents the ‘quantity’ of edges covered by the subgraphs. *C*\* is selected with optimal quality and quantity in terms of covering informative edges. λ_0_ can be tuned to either extract subgraphs with higher area density (i.e., low false-positive rates) or covering more high-weight edges using subgraphs with larger sizes (i.e., low false-negative rates). In general, *C*\* selection is robust for *τ*_0_ in the range of 0.4 to 0.7.

**S2 Fig.**
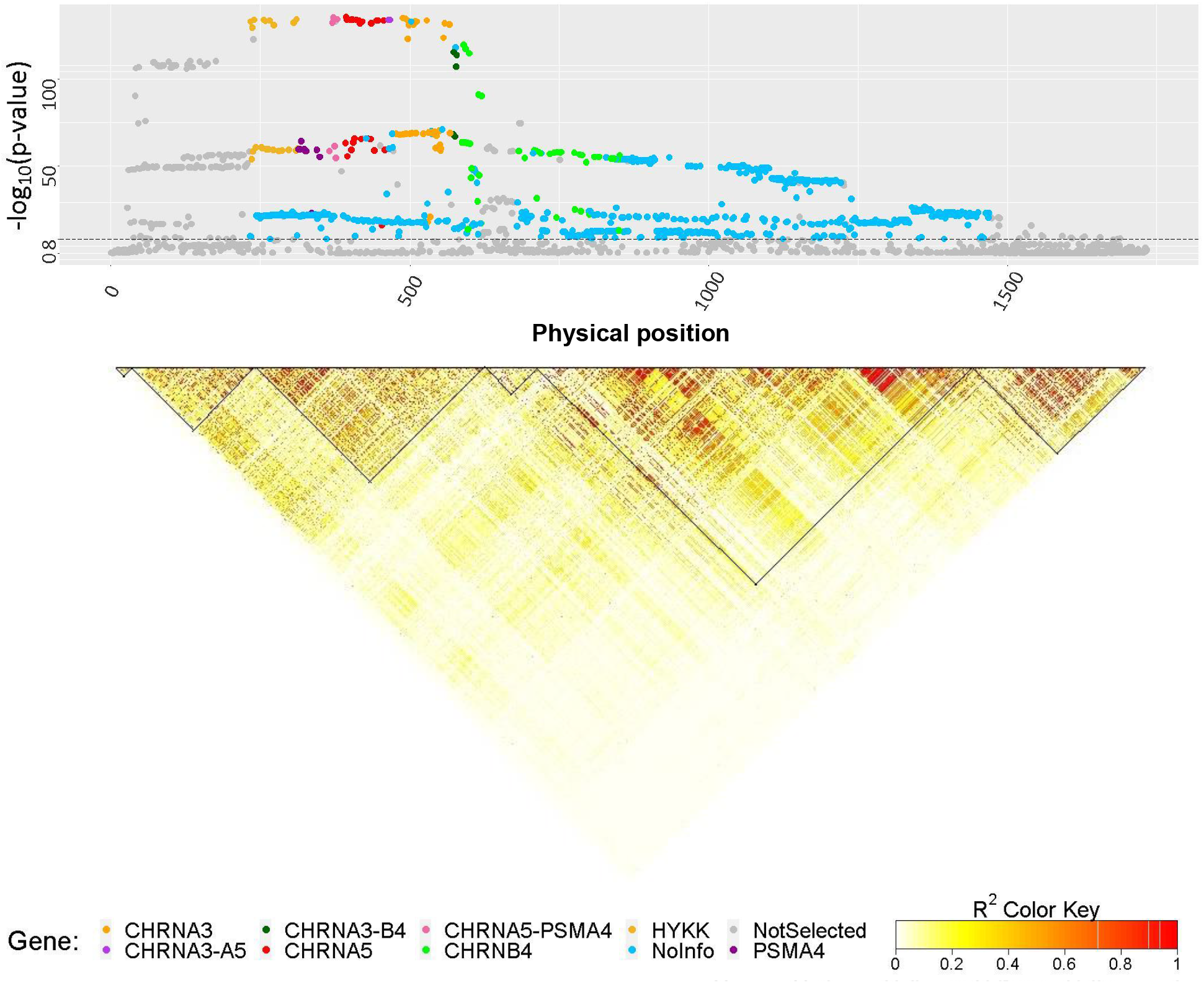
Comparison to the existing LD block detection method and corresponding fine-mapping results. We compare our dense LD block detection method with the cutting edge LD block detection method. The LD matrix in the figure shows the detected blocks by Big-LD [55]. Clearly, the eight Big-LD detected blocks are are much wider and sparse than the dense blocks from GSLD. Applying the Big-LD blocks to fused-LASSO, we select almost all SNPs regardless of the effect sizes, which may be misleading. The non-selected SNPs are colored in grey, and the selected SNPs are highlighted with corresponding genes (i.e., IREB2, CHRNA3, CHRNA5, CHRNB4, HYKK, and PSMA4).

## References

1. Klein RJ, Zeiss C, Chew EY, Tsai JY, Sackler RS, Haynes C, et al. Complement factor H polymorphism in age-related macular degeneration. Science. 2005;308(5720):385–389.

2. Mills MC, Rahal C. A scientometric review of genome-wide association studies. Communications biology. 2019;2(1):1–11.

3. Tam V, Patel N, Turcotte M, Bossé Y, Paré G, Meyre D. Benefits and limitations of genome-wide association studies. Nature Reviews Genetics. 2019;20(8):467–484.

4. Schaid DJ, Chen W, Larson NB. From genome-wide associations to candidate causal variants by statistical fine-mapping. Nature Reviews Genetics. 2018;19(8):491–504.

5. Visscher PM, Brown MA, McCarthy MI, Yang J. Five years of GWAS discovery. The American Journal of Human Genetics. 2012;90(1):7–24.

6. Visscher PM, Wray NR, Zhang Q, Sklar P, McCarthy MI, Brown MA, et al. 10 years of GWAS discovery: biology, function, and translation. The American Journal of Human Genetics. 2017;101(1):5–22.

7. Icick R, Forget B, Cloëz-Tayarani I, Pons S, Maskos U, Besson M. Genetic susceptibility to nicotine addiction: Advances and shortcomings in our understanding of the CHRNA5/A3/B4 gene cluster contribution. Neuropharmacology. 2020;177:108234.

8. Broekema R, Bakker O, Jonkers I. A practical view of fine-mapping and gene prioritization in the post-genome-wide association era. Open biology. 2020;10(1):190221.

9. Tibshirani R. Regression shrinkage and selection via the lasso. Journal of the Royal Statistical Society: Series B (Methodological). 1996;58(1):267–288.

10. Cho S, Kim H, Oh S, Kim K, Park T. Elastic-net regularization approaches for genome-wide association studies of rheumatoid arthritis. In: BMC proceedings. vol. 3. BioMed Central; 2009. p. 1–6.

11. Guan Y, Stephens M. Bayesian variable selection regression for genome-wide association studies and other large-scale problems. The Annals of Applied Statistics. 2011;p. 1780–1815.

12. Maller JB, McVean G, Byrnes J, Vukcevic D, Palin K, Su Z, et al. Bayesian refinement of association signals for 14 loci in 3 common diseases. Nature genetics. 2012;44(12):1294.

13. Dadaev T, Saunders EJ, Newcombe PJ, Anokian E, Leongamornlert DA, Brook MN, et al. Fine-mapping of prostate cancer susceptibility loci in a large meta-analysis identifies candidate causal variants. Nature communications. 2018;9(1):1–19.

14. He K, Kang J, Hong HG, Zhu J, Li Y, Lin H, et al. Covariance-insured screening. Computational statistics & data analysis. 2019;132:100–114.

15. Nicora G, Vitali F, Dagliati A, Geifman N, Bellazzi R. Integrated Multi-Omics Analyses in Oncology: A Review of Machine Learning Methods and Tools. Frontiers in Oncology. 2020;10:1030.

16. Wu C, Zhou F, Ren J, Li X, Jiang Y, Ma S. A selective review of multi-level omics data integration using variable selection. High-throughput. 2019;8(1):4.

17. Li C, Li H. Network-constrained regularization and variable selection for analysis of genomic data. Bioinformatics. 2008;24(9):1175–1182.

18. Jin J, Zhang CH, Zhang Q. Optimality of graphlet screening in high dimensional variable selection. The Journal of Machine Learning Research. 2014;15(1):2723–2772.

19. Wang X, Leng C. High-dimensional ordinary least-squares projection for screening variables. Journal of the Royal Statistical Society: Series B (Statistical Methodology). 2015;78(3):589–611.

20. Barrett JC, Fry B, Maller J, Daly MJ. Haploview: analysis and visualization of LD and haplotype maps. Bioinformatics. 2005;21(2):263–265.

21. Wang N, Akey JM, Zhang K, Chakraborty R, Jin L. Distribution of recombination crossovers and the origin of haplotype blocks: the interplay of population history, recombination, and mutation. The American Journal of Human Genetics. 2002;71(5):1227–1234.

22. Gabriel SB, Schaffner SF, Nguyen H, Moore JM, Roy J, Blumenstiel B, et al. The structure of haplotype blocks in the human genome. Science. 2002;296(5576):2225–2229.

23. Wu Q, Huang X, Culbreth A, Waltz J, Hong LE, Chen S. Extracting Brain Disease-Related Connectome Subgraphs by Adaptive Dense Subgraph Discovery. bioRxiv. 2020;.

24. Tibshirani R, Saunders M, Rosset S, Zhu J, Knight K. Sparsity and smoothness via the fused lasso. Journal of the Royal Statistical Society: Series B (Statistical Methodology). 2005;67(1):91–108.

25. Tibshirani RJ, Taylor J, et al. The solution path of the generalized lasso. The Annals of Statistics. 2011;39(3):1335–1371.

26. Arnold TB, Tibshirani RJ. Efficient implementations of the generalized lasso dual path algorithm. Journal of Computational and Graphical Statistics. 2016;25(1):1–27.

27. Friedman J, Hastie T, Höfling H, Tibshirani R, et al. Pathwise coordinate optimization. The annals of applied statistics. 2007;1(2):302–332.

28. Amos CI, Wu X, Broderick P, Gorlov IP, Gu J, Eisen T, et al. Genome-wide association scan of tag SNPs identifies a susceptibility locus for lung cancer at 15q25. 1. Nature genetics. 2008;40(5):616–622.

29. DeMeo DL, Mariani T, Bhattacharya S, Srisuma S, Lange C, Litonjua A, et al. Integration of genomic and genetic approaches implicates IREB2 as a COPD susceptibility gene. The American Journal of Human Genetics. 2009;85(4):493–502.

30. Marees AT, Gamazon ER, Gerring Z, Vorspan F, Fingal J, van den Brink W, et al. Post-GWAS analysis of six substance use traits improves the identification and functional interpretation of genetic risk loci. Drug and Alcohol Dependence. 2020;206:107703.

31. Erzurumluoglu AM, Liu M, Jackson VE, Barnes DR, Datta G, Melbourne CA, et al. Meta-analysis of up to 622,409 individuals identifies 40 novel smoking behaviour associated genetic loci. Molecular psychiatry. 2019;p. 1–18.

32. Friedman J, Hastie T, Tibshirani R. Regularization paths for generalized linear models via coordinate descent. Journal of statistical software. 2010;33(1):1.

33. Sudlow C, Gallacher J, Allen N, Beral V, Burton P, Danesh J, et al. UK biobank: an open access resource for identifying the causes of a wide range of complex diseases of middle and old age. Plos med. 2015;12(3):e1001779.

34. Chen S, Kang J, Xing Y, Zhao Y, Milton DK. Estimating large covariance matrix with network topology for high-dimensional biomedical data. Computational Statistics & Data Analysis. 2018;127:82–95.

35. Minicã CC, Mbarek H, Pool R, Dolan CV, Boomsma DI, Vink JM. Pathways to smoking behaviours: biological insights from the Tobacco and Genetics Consortium meta-analysis. Molecular psychiatry. 2017;22(1):82–88.

36. National Center for Biotechnology Information (NCBI) [Internet]. Bethesda (MD): National Library of Medicine (US), National Center for Biotechnology Information; 1988. Accessed: 2020-8-27. https://www.ncbi.nlm.nih.gov/.

37. Buniello A, MacArthur JAL, Cerezo M, Harris LW, Hayhurst J, Malangone C, et al. The NHGRI-EBI GWAS Catalog of published genome-wide association studies, targeted arrays and summary statistics 2019. Nucleic acids research. 2019;47(D1):D1005–D1012.

38. King DP, Paciga S, Pickering E, Benowitz NL, Bierut LJ, Conti DV, et al. Smoking cessation pharmacogenetics: analysis of varenicline and bupropion in placebo-controlled clinical trials. Neuropsychopharmacology. 2012;37(3):641–650.

39. Qiu W, Cho MH, Riley JH, Anderson WH, Singh D, Bakke P, et al. Genetics of sputum gene expression in chronic obstructive pulmonary disease. PloS one. 2011;6(9):e24395.

40. Yang L, Qiu F, Lu X, Huang D, Ma G, Guo Y, et al. Functional polymorphisms of CHRNA3 predict risks of chronic obstructive pulmonary disease and lung cancer in Chinese. PLoS One. 2012;7(10):e46071.

41. Berrettini W, Doyle G. The CHRNA5–A3–B4 gene cluster in nicotine addiction. Molecular psychiatry. 2012;17(9):856–866.

42. Joshi PK, Fischer K, Schraut KE, Campbell H, Esko T, Wilson JF. Variants near CHRNA3/5 and APOE have age-and sex-related effects on human lifespan. Nature communications. 2016;7(1):1–7.

43. Castaldi PJ, Cho MH, Zhou X, Qiu W, Mcgeachie M, Celli B, et al. Genetic control of gene expression at novel and established chronic obstructive pulmonary disease loci. Human molecular genetics. 2015;24(4):1200–1210.

44. Su Z, Marchini J, Donnelly P. HAPGEN2: simulation of multiple disease SNPs. Bioinformatics. 2011;27(16):2304–2305.

45. Purcell S, Neale B, Todd-Brown K, Thomas L, Ferreira MA, Bender D, et al. PLINK: a tool set for whole-genome association and population-based linkage analyses. The American journal of human genetics. 2007;81(3):559–575.

46. Newcombe PJ, Conti DV, Richardson S. JAM: a scalable Bayesian framework for joint analysis of marginal SNP effects. Genetic epidemiology. 2016;40(3):188–201.

47. Kichaev G, Yang WY, Lindstrom S, Hormozdiari F, Eskin E, Price AL, et al. Integrating functional data to prioritize causal variants in statistical fine-mapping studies. PLoS Genet. 2014;10(10):e1004722.

48. Kichaev G, Pasaniuc B. Leveraging functional-annotation data in trans-ethnic fine-mapping studies. The American Journal of Human Genetics. 2015;97(2):260–271.

49. Kichaev G, Roytman M, Johnson R, Eskin E, Lindstroem S, Kraft P, et al. Improved methods for multi-trait fine mapping of pleiotropic risk loci. Bioinformatics. 2017;33(2):248–255.

50. Goldberg AV. Finding a maximum density subgraph. University of California Berkeley; 1984.

51. Charikar M. Greedy approximation algorithms for finding dense components in a graph. In: International Workshop on Approximation Algorithms for Combinatorial Optimization. Springer; 2000. p. 84–95.

52. Tsourakakis CE. Mathematical and Algorithmic Analysis of Network and Biological Data. arXiv preprint arXiv:14070375. 2014;.

53. Stella XY, Shi J. Multiclass spectral clustering. In: null. IEEE; 2003. p. 313.

54. Bolla M. Spectral clustering and biclustering: Learning large graphs and contingency tables. John Wiley & Sons; 2013.

55. Kim SA, Cho CS, Kim SR, Bull SB, Yoo YJ. A new haplotype block detection method for dense genome sequencing data based on interval graph modeling of clusters of highly correlated SNPs. Bioinformatics. 2018;34(3):388–397.

